# Misregulation of the jasmonate signaling pathway leads to altered plant microbiota interaction and plant stress responses

**DOI:** 10.1101/2025.03.29.646076

**Authors:** Tung-Tse Lu, Miguelito Isip, Hung-Jui Sophia Shih, Silvina Perin, Ka-Wai Ma

## Abstract

The model plant *Arabidopsis thaliana* hosts diverse microbial communities collectively known as the microbiota. The plant microbiota is generally taxonomically structured. Some of the members can promote plant fitness including growth and stress tolerance. However, microbial imbalance can also result in deleterious effects, a phenomenon known as dysbiosis that was first coined in the gut microbiome field. To unveil the regulatory mechanism to maintain plant homeostatic interaction with microbiota, we performed screening using defined synthetic bacterial communities. We identified an Arabidopsis mutant with altered microbial profiles, an overall increase of microbial load and microbiota-dependent growth defects. Transcriptomic and chemical complementation analyses confirmed that the aforementioned microbiota-dependent phenotypes are contributed by an upregulation of the jasmonate signaling pathway. Upregulation of the jasmonate pathway further promotes microbial growth, possibly forming a positive feedback loop. Even though activation of the jasmonate signaling pathway is known to enhance plant stress tolerance, hyperactivation of the pathway alters plant tolerance or resistance against multiple stressors. Plant association with the microbiota together with proper regulation of the jasmonate signaling pathway are thus essential to maintain plant response to environmental stressors.

## Introduction

Plants are colonized by a multi-kingdom microbial community, generally known as the plant microbiota (Bulgarelli et al. 2013). Plant microbiota exhibit multifaceted impacts on their plant hosts. Some microbiota members are deleterious pathogens that exploit plant hosts under favorable conditions. Some are beneficial microbes that provide plant hosts with services ranging from growth promotion to facilitated nutrient uptake or stress resistance (Bulgarelli et al. 2013; Hassani et al. 2018). Plant root microbiota such as those of *Arabidopsis thaliana* is dominated by Proteobacteria, Actinobacteria and Bacteroidetes (Bulgarelli et al. 2012; Lundberg et al. 2012). Reconstitution experiments using synthetic communities comprising core microbiota members of microbial culture collection enables functional characterization of microbiota on plant hosts under controlled laboratory conditions (Zengler et al. 2019; Kremer et al. 2021; Ma et al. 2022). Such a reductionist approach is instrumental in shedding light on the roles of plant innate immunity (Ma et al. 2021; Teixeira et al. 2021), specialized metabolites chemistry (Huang et al. 2019; Vismans et al. 2022; Caddell et al. 2023; Thoenen et al. 2023; Yang et al. 2023; Jin et al. 2024; Liu et al. 2024; Su et al. 2024) and bacterial carbon preference in shaping plant microbiota composition (Schäfer et al. 2023). Although the plant microbiota, especially the bacterial microbiota, is taxonomically structured (Yeoh et al. 2017; Thiergart et al. 2020), it is subject to modulation due to environmental variations or still poorly understood host-mediated regulatory mechanisms. Given that microbiota activities and plant phenotypes are connected, further understanding of the principles governing plant-microbiota establishment can provide means to manipulate microbiota for enhancing agronomically important traits such as growth and yields without alteration in the plant genotypes.

The composition of host-associated microbiota sometimes deviates from the “norm”, a phenomenon that is coined dysbiosis. Dysbiosis was proposed in the field of the human gut microbiome to describe the microbial imbalance associated with diseases (Vangay et al. 2015; Levy et al. 2017). A major caveat to use this term stems from the implicit causal relationship between microbial imbalance and changes in host phenotypes, which is a classic circular argument that fails to distinguish dysbiosis as a cause, a consequence, or part of a feedback loop. Despite this unambiguity, understanding dysbiosis is crucial to provide a conceptual framework to account for the dynamics of microbiota and their consequences on plant host. Several dysbiosis mutants were reported in Arabidopsis. For example, mutations in the vesicle trafficking component *min7 and* three immune receptors/coreceptors including *fls2 efr cerk1* resulted in phyllosphere microbial imbalance. This quadruple mutant is characterized with a concomitant increase in the families Comamonadaceae, Xanthomonadaceae, Alcaligenaceae, Sphingomonadaceae and a decrease in Paenibacillaceae (Chen et al. 2020). Mutations in the membrane-bound NADPH oxidase *rbohD* (Pfeilmeier et al. 2021), the malectin-like domain-containing receptor *feronia* (Song et al. 2021) and the phytosulfokine receptor 1 *pskr1* (Song et al. 2023) are also associated with disease-like phenotypes. The disease-like phenotypes were attributed to an opportunistic *Xanthomonas* pathogen for *rbohD* (Pfeilmeier et al. 2021, 2024; Entila et al. 2024) and an overproliferation of *Pseudomonas* species for the others (Song et al. 2021, 2023). As such, yet unknown specificities regulated by the corresponding wild-type genes likely exert different impacts on the bacterial microbiota. Consistent with the previous reports showing that plant immunity plays a pivotal role coordinating plant-microbiota homeostasis (Lebeis et al. 2015; Ma et al. 2021; Teixeira et al. 2021), the dysbiosis mutants aforementioned are immunity-related. A recent genetic screen identified a S-acyltransferase mutant, *tip1,* exhibiting microbiota-dependent autoimmune phenotypes (Cheng et al. 2024). Together, these studies show a diversity of genes possibly involved in shaping microbiota composition.

We recently showed that individual members of the *A. thaliana* bacterial culture collection (Bai et al. 2015) exhibit contrasting abilities to modulate root immune responses (Ma et al. 2021). Using defined synthetic communities (SynComs) with taxonomically similar but distinct traits to interfere with root immune responses, we identified plant gene clusters showing differential response to these two groups of bacteria (Ma et al. 2021). We hypothesize that these differentially expressed gene clusters contain candidate genes important for healthy plant-microbiota interactions. To test this idea, we initiated a genetic screen to identify mutants with altered microbial communities, a *bona fide* feature of dysbiosis. We identified an Arabidopsis mutant, FLAG_264G01, with the T-DNA disrupting the second exon of a Class III peroxidase gene, *PEROXIDASE 5 (PER5)*. *per5* mutant exhibits phenotypes including microbial overgrowth and microbiota-dependent stunted growth. However, genetic analysis including the testing of independent CRISPR-edited *PER5* null mutants suggested that mutation of *PER5* alone is not sufficient to cause dysbiosis. Comparative transcriptomic analysis between wild type and *per5* hinted that the jasmonic acid (JA) pathway is upregulated in response to SynCom. JA and the derived methyl ester methyl jasmonate (MeJA), collectively known as jasmonates, are endogenous phytohormone well known for their regulatory role on plant growth and defenses (Howe et al. 2018). In Arabidopsis, jasmonates are primarily synthesized via the octadecanoid pathway in the plastids and promote the degradation of the repressor JAZ proteins through the E3 ubiquitin ligase COI1. JA pathway is upregulated in response to both abiotic and biotic stresses (Howe et al. 2018). However, the role of JA signaling regulating plant microbiota interaction is still unclear. Using two JA inducers, we directly demonstrate the upregulation of the JA pathway as a cause of the microbial imbalance phenotypes, supporting our claim that a feedback loop exists to shape plant microbiota composition. Upon perturbation of this feedback loop, plant microbiota interaction shifts and exert an impact on plant adaptation to environmental stressors.

## Results

### Identification of a mutant with microbiota-dependent stunted growth

To identify novel dysbiosis mutants, we performed reverse genetic screens using selected Arabidopsis T-DNA insertion mutants and a 16-member bacterial synthetic community SynCom (*At-*16SC1). The 16-member SynCom comprises members from 16 different bacterial families, representing a diversity of strains that were previously shown to exhibit robust colonization on Arabidopsis (Wippel et al. 2021). We prioritize our screening on mutants of genes with differential transcriptomic responses to microbiota of varying immunomodulatory traits (Ma et al. 2021). By focusing on mutants exhibiting microbiota-dependent phenotypes two weeks after inoculation compared to wild-type Ws4 plants (WT), we identified a mutant with significant stunted growth in the presence of SynComs (Fig 1a). Based on the FLAGdb/FST database (Samson et al. 2002), this mutant (FLAG_264G01) carries a T-DNA insertion at the *PEROXIDASE 5* gene (*PER5*). *PER5* was used as a defense marker gene (Zhou et al. 2020) but the function of *PER5* in pathogen resistance and regulation of microbiota is not reported. This mutant will be hereafter referred as *per5*.

**Fig. 1.**
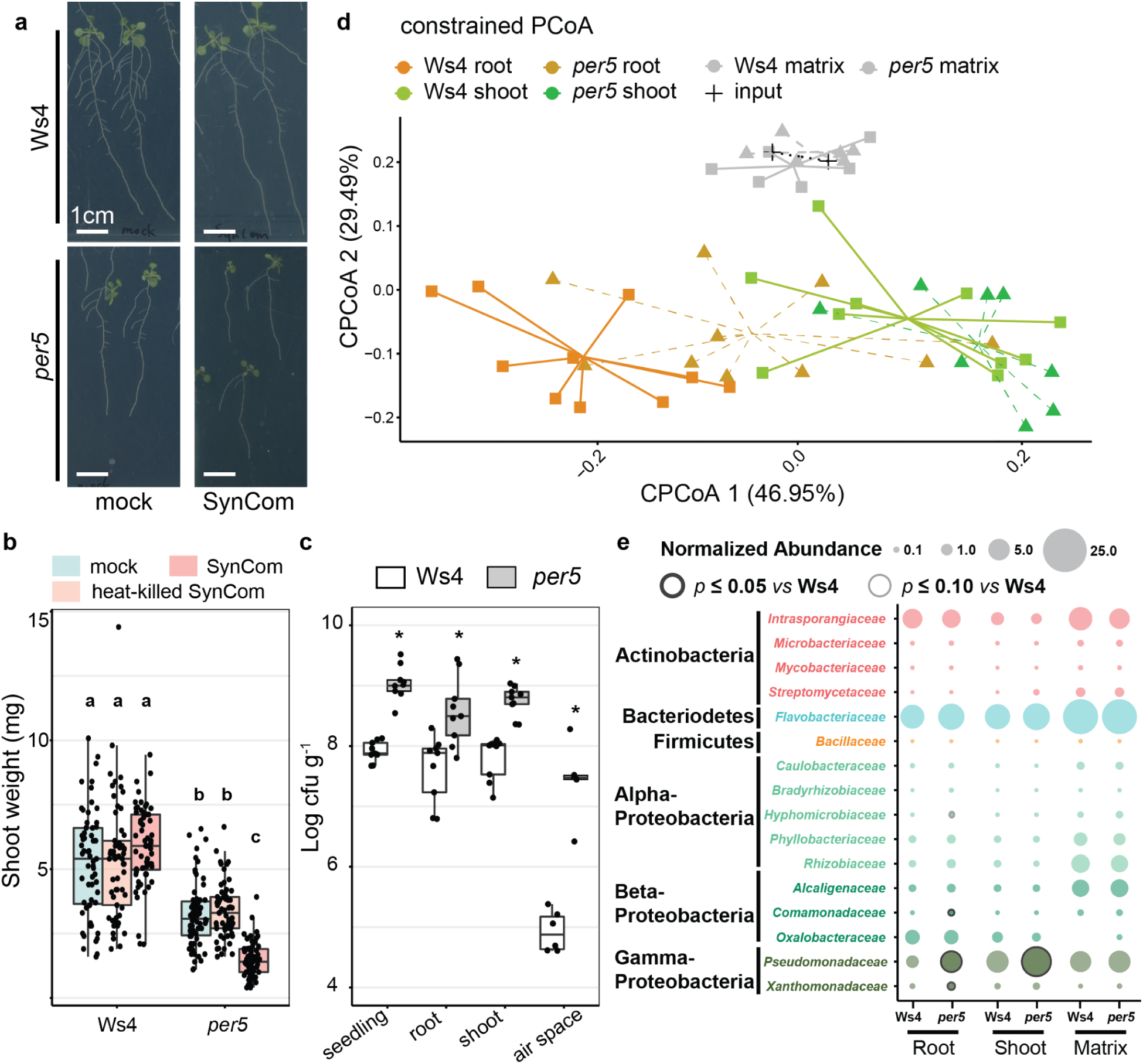
Identification of *per5* mutant with microbial imbalance phenotypes using synthetic bacterial communities. Wild-type Ws4 and *per5* were grown for two weeks on agar matrix in the presence of the 16-member SynCom *At*-16SC1. (a) Representative images of Ws4 and *per5* grown with *At*-16SC1. (b) Fresh shoot weight of plants treated with live or heat-killed SynCom. (c) Quantification of bacterial load in different compartments by live colony counts. (d) cPCoA plot of Bray-Curtis dissimilarities (ASV level) constrained by replicates. (e) Balloon plot showing the normalized abundance of individual families colonizing the indicated compartments and genotypes. Statistical significance was determined by Kruskal-Wallis followed by Dunn’s post-hoc test. Different letters and asterisks indicate statistical significance of *p*≤0.05 unless otherwise specified.

Under agar-grown axenic conditions with replete nutrients, *per5* mutants exhibited relatively mild growth defects with approximately 15-20% less fresh weight compared to the wild-type parent Wassilewskija accession (Ws-4) (Fig 1b, Supp Fig 1). While the presence of SynCom has a small growth promotion effect on WT, it resulted in up to 80% reduction of fresh weight for *per5* compared to the axenic control (Fig 1b, Supp). By contrast, inoculation with a heat-killed SynCom did not result in stunted growth (Fig 1b). *per5* showed more stunted growth when grown on agar plate compared to those grown in both potting and natural soils (Fig 1a, Supp Fig S1a), suggesting that the stunted growth phenotype is affected by the growing environment. To investigate whether microbiota-mediated growth inhibition *in per5* is dependent on the capacities of microbiota to activate or suppress immunity (Ma et al. 2021), we tested two additional SynComs: a five-member immune non-suppressive SynCom (NS1) and a five-member immune-suppressive SynCom (S1) (Ma et al. 2021). Inoculation of *per5* with these SynComs resulted in comparable growth inhibition to *At*-16SC1 (Fig 1b, Supp Fig S1b), suggesting that microbiota-mediated growth inhibition of *per5* is reproducible using three different SynComs. Occasionally, we observed chlorosis-like phenotypes on *per5* leaves (Fig 1b).

### *per5* growth phenotype is associated with microbial overgrowth and community shift

The growth defect of *per5* is associated with microbial overgrowth. Specifically, *per5* hosted approximately 10-fold higher total bacterial load compared to WT upon inoculation with the tested SynComs (Fig 1c, Supp Fig S1c). Unlike previously reported dysbiosis mutant that exhibits compartment-specific effect (Chen et al. 2020), microbial overgrowth is noted in both roots and shoots as well as the leaf air space, indicative of an overall change of microbiota (Fig 1c).

Next, we performed community profiling using partial sequences of the V5-V7 region of the *16S* ribosomal RNA gene. We spiked in a known amount of exogenous DNA during library preparation to enable normalization of the total microbial load. In line with our hypothesis that *per5* is a dysbiosis mutant with microbial imbalance, the microbial community of *per5* is significantly different from WT (Fig 1d). The community shift is observed in the plant-associated compartments i.e. shoot and root, but not in the agar matrix (Fig 1d). Using spike-in DNA for normalization, we showed that three Proteobacteria families including *Pseudomonaceae*, *Xanthomonadaceae* and *Comamonadaceae* are significantly enriched in *per5* roots compared to WT. Among these three families, *Pseudomonaceae* exhibits the highest enrichment from 1.23% to 4.54% (Fig 1e). We then performed a drop-out experiment to remove the *Pseudomonas* strain from the 16-member SynCom. The resultant 15-member SynCom still induced significant growth inhibition and enhanced total microbial load on *per5* but not WT (Fig 2a-b), suggesting that the deleterious effect from our 16-member SynCom is not due to the overproliferation of *Pseudomonaceae* strain alone.

**Fig 2.**
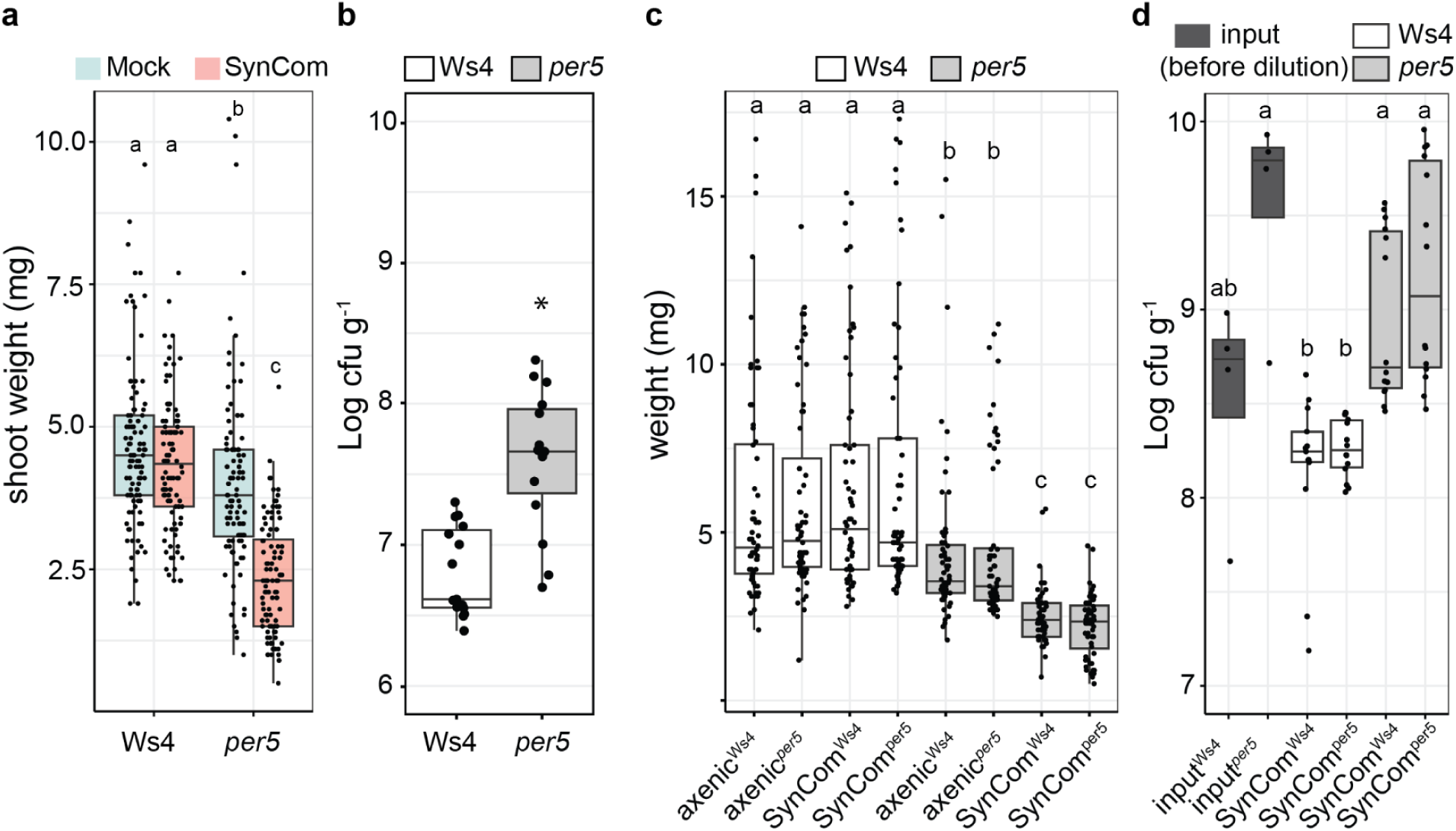
*per5* exhibits features that are atypical for dysbiosis mutants. Plants were grown for two weeks on agar matrix in the presence of the 15-member SynCom. The SynCom has the same composition of the *At*-16SC1 except for the drop out of the *Pseudomonas* strain R68. (a) and (b) Shoot weight and total microbial load of plants treated with the 15-member SynCom. (c) and (d) Shoot weight and total microbial load of plants inoculated with a Ws4 or *per5*-derived SynCom. Statistical significance was determined by Kruskal-Wallis followed by Dunn’s post-hoc test. Different letters and asterisks indicate statistical significance of *p*≤0.05 unless otherwise specified.

### *per5* exhibits atypical feature compared to known dysbiosis mutant

Following the definition of dysbiosis proposed by the human gut microbiome research, true dysbiosis should fulfil causality between specific microbiota composition and changes in host phenotypes. To address the role of microbial imbalance on plant performance, we prepared WT or *per5*-derived homogenate carrying the respective microbial communities as starting inocula, followed by inoculation on WT or *per5* plants. To account for the difference in microbial titer (Fig 2d), both derived SynComs are diluted to comparable microbial titers before inoculation. The plant growth phenotypes from these inoculation experiments exhibit more variation, possibly due to the interference by co-extracted plant factors including nutrients and differences in microbiota extraction efficiency. However, we conclude that *per5*-derived microbes cannot recapitulate microbiota-mediated plant growth inhibition and microbial overgrowth in wild-type Ws4 plants (Fig 2c-2d). Therefore, *per5* exhibits characters that are different from classic dysbiosis mutants. Together, we identified a mutant with microbiota-dependent stunted growth, shift in microbiota composition and enhanced microbial load.

### *per5* loss-of-function is not sufficient to cause dysbiosis

*per5* mutant (FLAG_264G01) was annotated to have a T-DNA insertion at the second exon of the class III peroxidase gene *PER5* (Supp S2a). There are 73 Class III peroxidases in *A. thaliana*, implicated in functions ranging from plant development, abiotic and biotic stress responses (Shigeto and Tsutsumi 2016). Consistent with a putative role of PER5 in modulating reactive oxygen species (ROS) level, *per5* mutant has reduced basal and microbe-induced ROS as shown by DAB staining (Supp S2b). Although *PER5* carries a T-DNA insertion at the exon, expression analyses using primers targeting both upstream (R1) and downstream regions (R2 and R3) of the *per5* T-DNA insertion site showed that *per5* transcripts were expressed (Supp S2c). To unambiguously investigate the contribution of *PER5* loss-of-function to the dysbiosis-like phenotypes, we generated two independent CRISPR deletion mutants of *PER5*, *per5 #3-1* and *per5 #5-2* CRISPR in the Col-0 background (Supp S2d). None of them showed any microbiota-dependent growth inhibition phenotypes (Supp S2e).

We expanded our analyses on the genes surrounding *PER5* locus. The expressions of two neighboring genes within 4 kb proximity of the *PER5* locus, another peroxidase gene *PEROXIDASE 4 (PER4)* and the mitochondrial CoA transporter (*CoAC1)*, were also affected in *per5*. To assess the contribution of these two genes, *PER4* and *CoAC1,* to the phenotypes, we tested a knockdown mutant of *per4* (*prx4-1*), a hypermorphic mutant of *PER4* (*prx4-2*) (Arnaud et al. 2017) and two null mutants of *CoAC1 (coac1-1* and *coac1-2)*. None of them showed any microbiota-dependent stunted growth (Supp S2e). Therefore, loss-of-function of *per5* alone is not sufficient to cause dysbiosis. However, we cannot exclude the possibility that the phenotypes are contributed by unknown epistatic interactions involving multiple genes or still uncharacterized polymorphisms in the *per5* background. The actual causative factor(s) contributing to microbial imbalance requires further investigation.

### The JA pathway is upregulated in the *per5* mutant

Although the genetic determinant(s) contributing to microbial imbalance is still unclear, we sought to identify biological processes associated with our phenotypes by performing transcriptomic analyses on WT and *per5* in the presence or absence of the 16-member SynCom. Using principal component analysis (PCA), the first PC accounts for 73% of the variation, separating root and shoot samples. The second PC separates axenic from SynCom-treated plants (4% variation). Although plant genotype (e.g. Ws4 vs *per5*) only explains a minority of the variation, *per5* still form distinct clusters from WT under both axenic and SynCom-inoculated conditions in both roots and shoots (Fig 3a). The more pronounced separation of transcriptomes in the presence of SynCom compared to axenic plants is consistent with the plant phenotypes e.g. more stunted growth in the presence of SynCom (Fig 1a-b).

**Fig 3.**
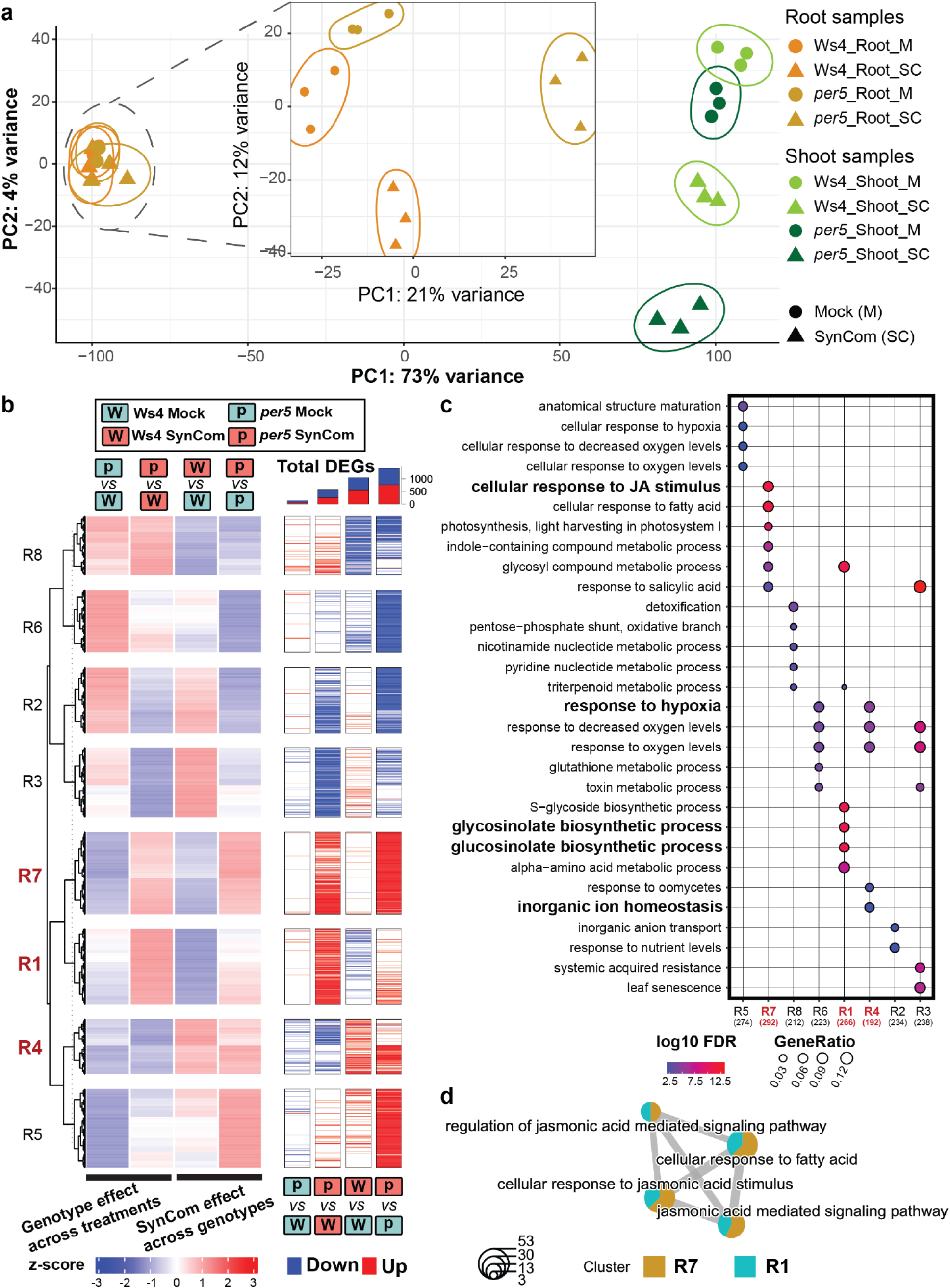
*per5* has distinct transcriptomic responses to the synthetic bacterial community. Plants were grown for two weeks on agar matrix in the presence of the SynCom *At*-16SC1. Root and shoot samples were harvested separately and subjected to RNA-seq analysis. (a) Principal component analyses showing the separation of transcriptomes in different samples. The insert corresponds to the result using root-associated samples only. (b) Heatmaps summarizing the expressions of genes across both genotypes in roots. DEGs were calculated by pairwise comparisons (log2FC>1.5, *p*<0.05). (c) Top five significantly enriched GO terms associated with each cluster. (d) GO terms related to JA processes and their association with clusters R1 and R7. Size of pie charts corresponds to the number of DEGs.

To better visualize the genotype-specific differences, we performed hierarchical clustering separately on z-score transformed root and shoot dataset. We identified sets of differentially expressed genes (DEGs, log2FC>1.5, *p*<0.05) using pairwise comparisons (Fig 3b), followed by GO term enrichment analysis (Fig 3b, Supp S3). As such, we group our dataset into eight clusters. From our root dataset, we identified a SynCom-responsive cluster, **R4**, which is upregulated across both genotypes (Fig 3c). GO term enrichment suggests that cluster **R4** is linked with hypoxia and inorganic ion homeostasis. In line with previous report (Ma et al. 2021), these biological processes represent GO terms generally responsive to bacterial SynComs. We identified another SynCom-responsive cluster, **R1**, exhibiting contrasting responses across genotypes e.g. upregulated in *per5* but downregulated in WT. **R1** genes are associated with functions related to glucosinolate biosynthesis and regulation of the JA signaling pathway (Fig 3c-3d). We also identified cluster **R7**, another SynCom-responsive cluster, which is more strongly upregulated in *per5* but not WT. **R7** is enriched with genes related to defense and JA responses (Fig 3c-3d). Although JA signaling is known to affect plant growth and defense responses, the role of JA on plant-microbiota interaction is relatively unclear (Doornbos et al. 2011; Carvalhais et al. 2013; Liu et al. 2017; Berendsen et al. 2018). We thus focus on the JA pathway for further analyses.

To investigate whether JA-related processes are affected at the biosynthesis or the signaling level, we looked at the expression levels of genes involved in JA biosynthesis (e.g. *LOX3*, *LOX4*), regulation (e.g. multiple JAZs) and key transcription factors/markers (e.g. *PDF1.2*, *MYC2*, *VSP1*). All of them all upregulated in *per5* compared to WT in the presence of SynCom (Fig 4a, Supp S4), suggesting the JA signaling pathway is upregulated. To our surprise, multiple JAZs are also upregulated. JAZs are repressor proteins that negatively regulate JA signaling (Howe et al. 2018). We speculate that upregulations of JAZs are the result of negative feedback in response to hyperactivation of the JA pathway in the *per5* background.

**Fig 4.**
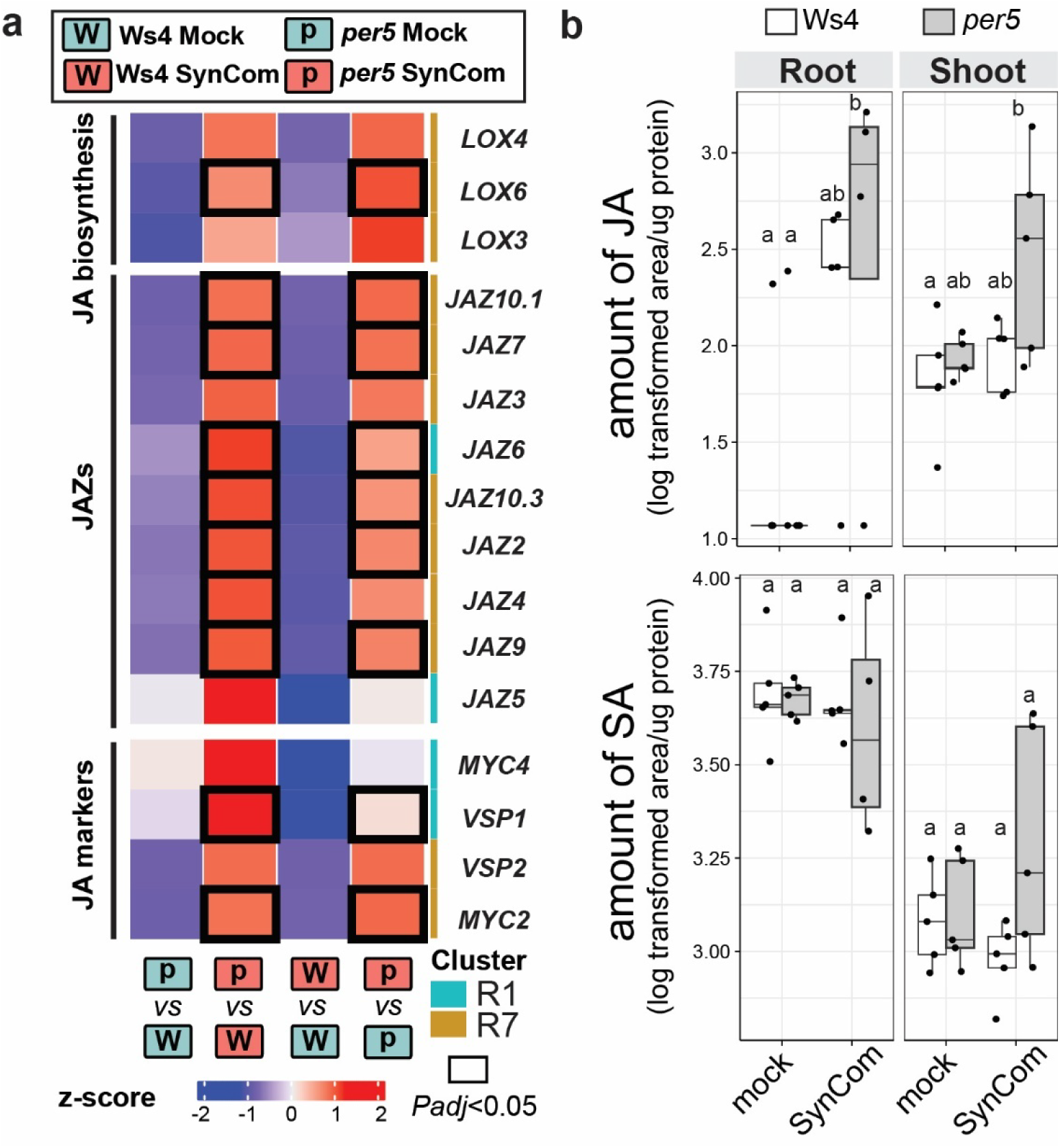
JA levels and related genes are upregulated in *per5* in response to the synthetic community. (a) Normalized relative expression levels of selected genes involved in JA-related processes in roots (related to Fig 3b). Genes of statistical significance (log2FC>1.5, *p*<0.05) are indicated with blocked lines. (b) The amount of JA and SA of two weeks old Ws4 and *per5* plants in the presence or absence of SynCom *At*– 16SC1. Statistical significance was determined by Kruskal-Wallis followed by Dunn’s post-hoc test. Different letters indicate statistical significance of *p*≤0.05 unless otherwise specified.

To understand the dynamics of JA levels in response to bacterial SynCom, we also quantify the levels of JA across both genotypes. We could barely detect JA in axenic roots. However, inoculation with SynCom *At-*16SC1 results in elevated levels of JA. The increases are more significant in *per5* than WT (Fig 4b, Supp S5). By contrast, there is no significant increase of another phytohormone, e.g. salicylic acid (SA), suggesting that the upregulation of JA cannot be explained by antagonism between JA and SA pathway.

### Enhanced JA signaling contributes to microbial imbalance

If microbiota-mediated upregulation of JA indeed contributes to the microbial imbalance phenotypes, we expect that exogenous application of JA inducers in the presence of SynCom can phenocopy *per5* phenotypes on WT. To differentiate between direct chemical-mediated growth inhibition and growth modulation through interaction with the microbiota, we tested different concentrations of methyl-jasmonate (MeJA) and 2,6-dichlorobenzonitrile (DCB). DCB is a cell wall synthesis inhibitor known to induce JA signaling (Ellis et al. 2002b). First, we identified the highest concentrations of MeJA and DCB that do not lead to significant plant growth inhibition (Fig 5a). Exogenous application of these two chemicals under non-inhibitory concentrations together with the SynCom reproduces the growth inhibition phenotypes in wild-type plants (Fig 5b), suggesting that upregulation of the JA pathway results in microbiota-dependent growth inhibition.

**Fig 5.**
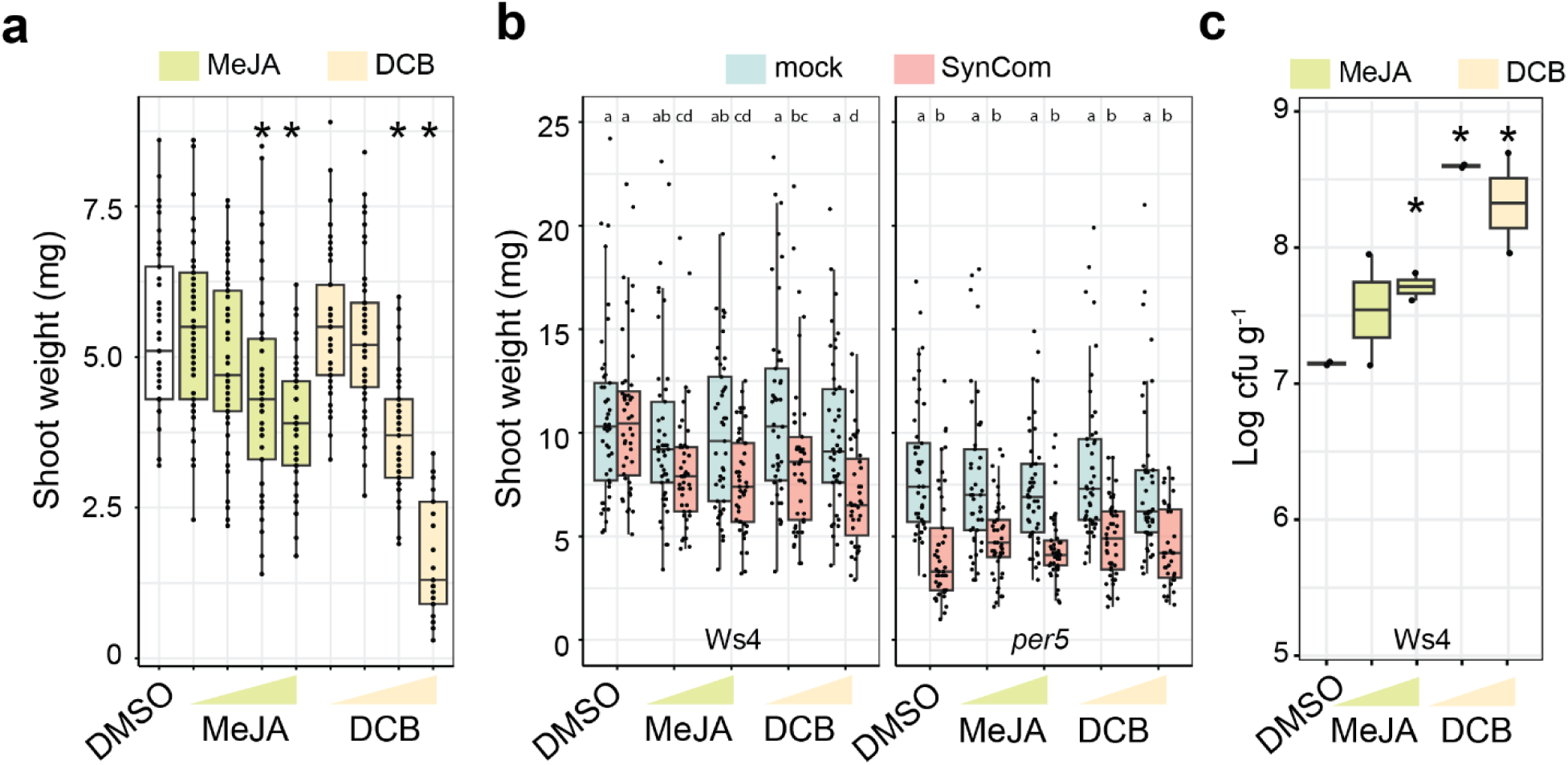
Treatment with two JA inducers reproduces *per5* phenotypes on WT. (a) Effect of different concentrations of MeJA and DCB on plant growth. MeJA: 0.1, 0.25, 0.5, 1.0µM; DCB: 0.05, 0.1, 0.25, 0.5µM. Note that the non-inhibitory concentration of MeJA (0.1 and 0.25 µM) and DCB (0.05 and 0.1 µM) are used in the subsequent experiment with SynCom. (b) Treatment with different concentrations of MeJA or DCB induced SynCom-dependent growth inhibition on WT. MeJA: 0.1, 0.25µM; DCB: 0.05, 0.1µM. (c) Treatment with different concentrations of MeJA or DCB increased the total microbial load on wild-type Ws4 plants. MeJA: 0.5, 2.5 uM; DCB: 0.1, 0.5 uM. Statistical significance was determined by ANOVA. Different letters and asterisks indicate statistical significance of *p*≤0.05 compared to the DMSO control unless otherwise specified.

Next, we tested whether exogenous applications of MeJA and DCB result in microbial over-proliferation. Using a similar setup and a 16-member SynCom, *At-*16SC2, we showed that MeJA and DCB treatment led to 5-10-fold higher microbial loads on wild-type plants (Fig 5c). However, the threshold required to cause growth inhibition and microbial overgrowth seem to differ. Since microbial overgrowth is coupled with stunted growth under our experimental conditions, we cannot distinguish between primary and secondary effects upon upregulation of the JA pathway, e.g. plant morphological changes, on microbiota colonization. Together, we provide evidence that plant microbiota composition and the plant JA signaling pathway is regulated by an unknown feedback mechanism, possibly affected by yet unidentified genetic determinant(s) in the *per5* mutant background.

### Misregulation of the jasmonate pathway in *per5* alters plant responses to multiple stressors

Recently, the role of microbiota in autoimmunity has been reported (Cheng et al. 2024). We thus asked whether *per5* has heightened immunity level. Autoimmune mutants are characterized by higher expressions of defense marker genes. We selected four defense marker genes such as *SARD1*, *CBP60g, FRK1* and *PR1* for further analyses (Wang et al. 2011). While *SARD1, CBP60g* and *FRK1* are upregulated in response to SynCom across both genotypes in the shoot, the expression of *PR1* appears to be upregulated only in the *per5* mutant (Fig 6a). To directly investigate whether *per5* is more resistant to bacterial phytopathogens, we syringe infiltrate *per5* and WT plants with the hemibiotrophic pathogen *Pseudomonas syringae pv. tomato PtoDC3000*. The pathogen load of *per5* is significantly lower than WT (Fig 6b), suggesting that *per5* is more resistant to *Pseudomonas* pathogen. Next, we tested whether upregulation of JA in *per5* is linked with altered plant tolerance to an abiotic stressor such as high salinity. We observed that *per5* mutant is hypersensitive to salt compared to WT (Fig 6c). Together, these results indicate that *per5* mutant has altered resistance and tolerance to multiple stressors, possibly linked to a misregulation of the jasmonate signaling pathway.

**Fig 6.**
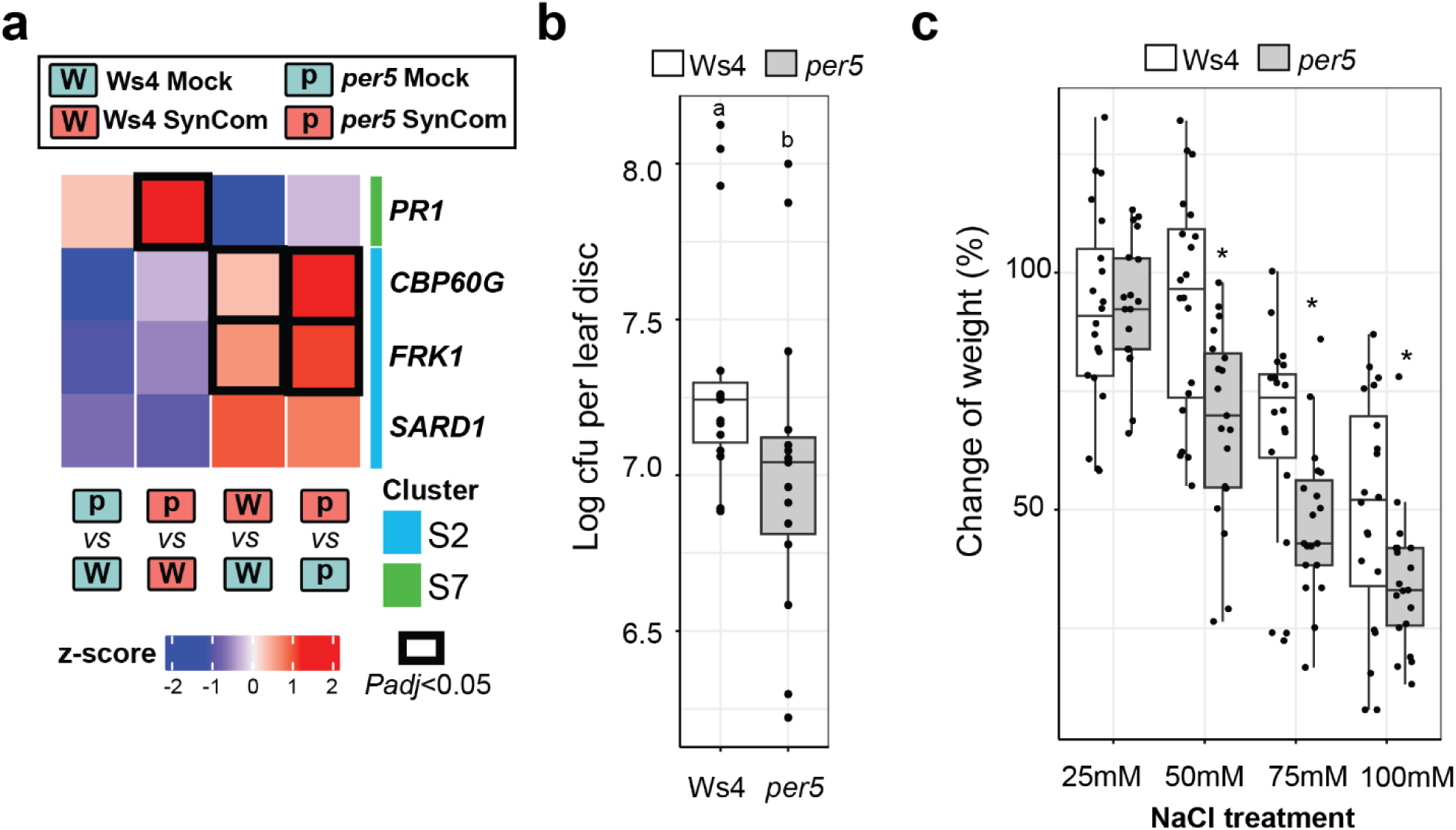
*per5* mutant has altered responses to multiple stressors. (a) Normalized relative expression levels of selected genes involved in defense responses in shoot. Genes of statistical significance (log2FC>1.5, *p*<0.05) are indicated with blocked lines. (b) Bacterial load of the foliar bacterial pathogen *Pto*DC3000 on WT and *per5* plants. Bacterial load was quantified based on colony forming unit (cfu) per unit area of leaf disc. (c) Sensitivity of plants with an increasing concentration of salt. Sensitivity was expressed in terms of percentage change of fresh weight compared to the mock treatment across both genotypes. Note that *per5* but not WT shows reduced fresh weight at 50mM salt treatment.

## Discussion

### Identification of *per5* as a mutant with microbial imbalance

Over the last decade, multiple factors implicated in plant microbiota establishment have been identified (Bulgarelli et al. 2012, 2013; Lundberg et al. 2012; Edwards et al. 2015; Wagner et al. 2016; Ma et al. 2021; Schäfer et al. 2023; Entila et al. 2024). In the case of root microbiota, edaphic factors play deterministic roles by shaping the composition of the initial soil inoculum for subsequent root colonization. Cell wall features, root exudation linked with microbial resources preferences, and plant innate immunity work together to fine-tune microbiota composition. Despite these advances, the genetic basis underlying plant microbiota homeostasis remains unclear. Several dysbiosis mutants were reported sharing a common or suspected role related to plant innate immunity (Xin et al. 2016; Chen et al. 2020; Pfeilmeier et al. 2021; Song et al. 2021, 2023; Cheng et al. 2024). In this study, we reported a novel *per5* mutant with microbial imbalance phenotypes. *per5* exhibits *bona fide* features of dysbiosis but is different in some ways. First, *per5*-derived microbiota failed to reproduce the dysbiosis phenotypes on wild type plants, suggesting a mechanism to restore the otherwise “abnormal” *per5*-derived community back to normal. Second, the associated microbial overproliferation is not specific to a single bacterial family. Third, although *per5* resembles autoimmune mutants such as *snc1* (Zhang et al. 2003) and *tip1* (Cheng et al. 2024) with a stunted growth phenotype, they differ in several ways. *snc1* and *tip1* exhibit strong growth defects under axenic conditions and the presence of microbes alleviates this phenotype. In contrast, microbiota aggravates the stunted growth phenotype in *per5* (Fig 1a). Stunted growth of *per5* is unlikely the result of elicitor hypersensitivity as treatment of *per5* with the heat-killed SynCom, presumably carrying a cocktail of heat-stable elicitors, did not result in any obvious stunted growth (Fig 1b).

### Upregulation of the JA pathway contributes to altered plant microbiota interaction

Based on transcriptome analyses, we showed that the JA pathway is upregulated in *per5* in the presence of microbiota (Fig 3-4). The role of JA in shaping microbiota composition has been investigated before, but their conclusions vary. While one study showed that JA treatment led to rhizosphere community shifts in *Arabidopsis* (Carvalhais et al. 2013), other studies suggested the otherwise (Doornbos et al. 2011; Liu et al. 2017; Berendsen et al. 2018). Such discrepancies are likely compounded by variations in soil parameters, microbiota composition (Doornbos et al. 2011) and the method of community analyses used (e.g. sequencing versus PhyloChip). Additionally, volatiles such as MeJA can act on microbes by affecting biofilm formation, thus contributing to a community shift (Kulkarni et al. 2024). To directly investigate the possibility of a feedback loop between JA and the bacterial microbiota, we performed reconstitution experiment using SynCom. Colonization by SynCom results in higher levels of JA. Conversely, treatment of two JA inducers reproduces *per5* phenotypes including microbiota-dependent stunted growth and microbial overgrowth (Fig 5), suggesting that the upregulation of JA signaling can directly cause changes in plant-microbiota interactions. We speculate that this JA-dependent feedback loop functions within physiological relevant ranges in wild-type plants upon association with the SynCom. However, due to unidentified genetic perturbation in the *per5* mutant, this feedback loop is amplified. The resultant upregulation of the JA pathway does not only compromise plant growth but also impact plant responses to both abiotic and biotic stressors (Fig 6). Together, our results underscore the importance for proper regulation of the JA pathway in maintaining healthy plant microbiota interaction. However, the genetic determinant(s) and mechanism(s) leading to altered plant microbiota interaction in the *per5* background requires further investigation.

Upregulation of JA signaling is essential for plant adaptation to environmental stresses. Upregulation of the JA pathway can promote plant disease susceptibility through the suppression of the SA branch, a virulence strategy deployed by some pathogenic strains (Cui et al. 2005; Zheng et al. 2012). However, enhanced pathogen resistance has also been reported to be associated with higher JA signaling as in the *cev* mutant (Ellis and Turner 2001; Ellis et al. 2002a), corresponding to a mutation in the cellulose synthase CeSA3 (Ellis et al. 2002b). In line with this, treatment of plants with the cell wall inhibitor DCB also promotes microbial colonization (Fig. 5c). Defect in cell wall integrity can activate plant innate immunity (Zhai et al. 2024). It is thus possible that the enhanced resistance against *Pto*DC3000 of *per5* is due to altered immunity after microbiota colonization. Additionally, community shift or microbial overproliferation may contribute to pathogen resistance through interbacterial competition or activation of induced systemic resistance (ISR) (Pieterse et al. 1998; Pozo et al. 2008; Van der Ent et al. 2008; Zamioudis et al. 2014).

In addition to hemibiotrophic pathogens, upregulation of JA signaling is also known to confer abiotic stress tolerance. For example, in response to high salinity, JA is synthesized and results in the degradation of JAZ repressor proteins through the COI1-dependent proteasomal degradation pathway. Degradation of JAZ8 leads to the derepression of the salt-responsive NF-YA1-YB2-YC9 complex (Li et al. 2024), thereby contributing to salt tolerance. Enhanced level of JA is often associated with the degradation of JAZs (Howe et al. 2018). However, in the case of *per5*, enhanced JA level is coupled with the transcriptional upregulation of multiple *JAZ*, including *JAZ8* in the shoot (Fig 4, Supp S4). Upregulation of JAZs may possibly account for the reduced salt stress tolerance in *per5* mutants despite an overall upregulation of the JA signaling pathway. Together, these results highlight the complexity of the underlying regulatory network. Possible interaction with the microbiota should be considered, with the overarching goal of promoting both plant growth and stress tolerance.

## Materials and methods

### Plant Materials and growth conditions

*Arabidopsis thaliana* ecotype Columbia (Col-0, CS60000), Wassilewskija (*Ws4)* were lab stocks. *per5* (FLAG_264G01), *PER4* knock-down mutant (SALK_110617C, *prx4-1*), PER4 hypermorphic mutant (SALK_044730C, *prx4-2*) (Arnaud et al. 2017) and *COAC1* null mutants (SALK_091671C, *coac1-1*; SALK_087365C, *coac1-2*) were obtained from the Nottingham Arabidopsis Stock Center (NASC). *Arabidopsis* seeds were surface-sterilized according to a published protocol (Ma et al. 2022). Briefly, seeds were incubated 5 minutes each with 70% ethanol twice, followed by a brief wash with 100% ethanol. Seeds were washed thoroughly with sterile water three times and cold-stratified seeds were sowed on agar plates (1%, Difco Bacto Agar, BD Biosciences) supplemented with half-strength Murashige and Skoog (MS) medium (M0222, Duchefa, NL), 0.1g/L 2-(*N*-morpholino)ethanesulfonic acid (MES, pH 5.7). Plants were grown in a growth chamber under short-day conditions (10 hr light, 14 hr dark) at 21°C/19°C cycle, 65% relative humidity and light intensity of 120 mE m^−2^ sec^−1^ or a controlled growth room.

### gRNA design and the generation of *PER5* CRISPR lines

4 pairs of gRNA sequences targeting PER5 gene were designed using the default settings of the ChopChop program (Labun et al. 2019). Hybridized gRNAs were inserted into the recipient vector pDEG347 though shuttle vectors using the published restriction-ligation protocol (Stuttmann et al. 2021). Arabidopsis was transformed by Agrobacterium GV3101 pMP90 carrying the final transformation vector using the floral dipping method. Briefly, inflorescences of plants were dipped into an Agrobacterium solution (a LB-grown saturated bacterial culture resuspended in 5% sucrose solution and 0.02% Silwet L-77). Primary transformants were selected by resistance to the herbicide phosphinothricin. Oligonucleotides corresponding to the target sites can be shared upon request.

### Culture condition for bacteria

Culture collection members, Agrobacterium GV3101 and *Pseudomonas syringae Pto*DC3000 were grown on 50% tryptic soy broth (TSB) agar plate (Sigma-Aldrich, USA or cyrusbioscience, Taiwan), Luria-Bertani broth or King’s B medium (HiMedia, USA) at 25 °C or 28 °C from one to four days. For the preparation of SynCom, fully grown bacterial cultures were pelleted by centrifugation at 8k g for 5 min, followed by two washes and resuspension in 10mM MgSO_4_. Strains were inoculated to warm agar medium at a concentration of OD_600_=0.0005 for each strain. Two 16-member SynComs (*At-*16SC1 and *At*-16SC2) were prepared. The composition of *At*-16SC1 is similar to the previously published *At*-SC4 (Wippel et al. 2021) except for the replacement of the Betaproteobacteria strain root1221 with stain29. *At*-16SC2 is similar to *At*-16SC1 except for the replacement of strain R418 by R336D2 (both of them are Oxalobacteraceae strains) due to the loss of R418 after long-term storage.

### Reinoculation experiment

Ws4 and *per5* plants were grown with *At-*16SC1 for two weeks. Ws4 and *per5* seedlings were harvested and homogenized in 10mM MgSO_4_. The inocula were diluted to final concentrations of 0.01g/ml and 0.001g/ml for Ws4 and *per5*, respectively. Similar bacterial titers were validated by plate counting method. The plant extracts were mixed with full strength MS agar to obtain half strength MS medium (final concentration of MES and agar remain unchanged).

### DAB staining

7-day Ws4 or *per5* seedlings with or without *Pseudomonas* strain root9 were stained with 1mg/ml DAB prepared in a 10mM phosphate buffer (pH7.0) for 1 hr. After three washes, the seedlings were observed under a stereomicroscope. The intensity of DAB at the root tip was quantified by ImageJ.

### Quantification of JA and SA levels

Two weeks old Ws4 and *per5* plants were germinated on half strength MS agar plate supplemented with SynCom *At*-16SC1. Plants are separated from the growth media by a 100 micron nylon mesh (Nitex Cat# 03-100/44, Sefar). Roots and shoots were harvested separately. Around 30 mg of tissues were pulverized (Tissuelyser II, Qiagen). Metabolites were extracted using an extraction buffer (ACN:MeOH:0.06%HCL 40:40:20 (v:v:v) + 20 ng/mL salicylic d4 as an internal standard). Briefly, 1ml pre-cooled (−20°C) extraction buffer was added to each sample, followed by incubation at 1,500 rpm in a thermomixer for 30 min at 4 °C. Samples were centrifuged at 21,000g for 10 min. The pellets were used to determine protein concentrations using a bicinchoninic acid assay (BCA). The cleared supernatants were transferred to a new tube and dried in a Speed Vac concentrator (Centrivap; Labcono) at 10°C at 1,000 rpm. Dried samples were re-suspended in 100 μl solution containing ACN (LC-MS grade): NH4 for (10mM) + FA 0,15% (50:50 (v:v)). Samples were vortexed for 30s and centrifuged for 2 min at 10,000g and 4 °C. The cleared supernatants were transferred to 200 μl glass inserts (CZT; Germany). All samples were placed in an Acquity iClass UPLC (Waters) sample manager held at 10 °C. The UPLC was connected to an Orbitrap, equipped with a heated ESI (HESI) source (QExactive, Thermo Fisher Scientific). 2 μl was injected onto a 100 × 1.0 mm HSS T3 C18 UPLC column, packed with 1.8 μm particles (Waters). The flow rate of the UPLC was set to 100 μl min^−1^ and the buffer system consisted of buffer A (10 mM ammonium Formate and 0.15% Formic acid in UPLC-grade water) and buffer B (UPLC-grade acetonitrile). The UPLC gradient was as follows: 0–1 min 90% A, 1–8 min 90–10% A, 8–10 min 10% A, 10.01-12 min 90% A. This leads to a total runtime of 12 min per sample.

The QExactive mass spectrometer was operating in negative ionization mode scanning a mass range between m/z 50 and 750. The maximal ion time was set to 200 ms, and the HESI source was operating with a spray voltage of 2.75 kV in negative ionization mode. The ion tube transfer capillary temperature was 350 °C, the sheath gas flow 50 arbitrary units (AU), the auxiliary gas flow 14 AU and the sweep gas flow was set to 3 AU at 380 °C. All samples were analyzed in a randomized run order. Targeted data analysis was performed using the Quan module of the TraceFinder 4.1 software (Thermo Fisher Scientific) in combination with a sample-specific compound database, derived from measurements of commercial reference compounds (Sigma). Samples below the detection limit are not excluded from the analyses. Instead, these samples are replaced by half of the lowest detection values (DL/2).

### 16S amplicon sequencing and community profiling

Libraries were processed according to previously published protocol (Ma et al. 2021). Briefly, plant roots and shoots were harvested separately unless otherwise specified. Total DNA was extracted using the FastDNA SPIN Kit for Soil (MP Biomedicals) according to the manufacturer’s instructions. Samples were diluted to 3.5 ng/μl before being used as templates in a three-step protocol published previously (Ma et al. 2021). Spike-in DNA (Ordon et al. 2024) was added during the first round PCR reaction to enable normalization of the data. Forward and reverse sequencing reads were denoised and demultiplexed separately according to the barcode sequences using QIIME (Caporaso et al. 2010) with the following parameters: *phred*=30; *bc_err*=2. After quality-filtering, merging of paired-end reads, amplicon tags were then aligned to the reference sequences using USEARCH (*uparse_ref* command) (Edgar 2010). Correction of PCR and sequencing errors was performed using the rbec pipeline (Zhang et al. 2021). An amplicon sequence variant (ASV) table for each strain was generated. This table was used for subsequent diversity analyses. Bray-Curtis dissimilarity index was calculated using the *vegdist* function in the vegan package (Oksanen et al. 2025). Constrained PcoA was performed using the vegan capscale function on the Bray-Curtis dissimilarity matrices, constraining by the following formula.

**genotype * compartment + Condition(biological.replicate * technical.replicate)**

One out of three biological replicates shown Fig. 1c was excluded from analysis due to contamination issue for the reconstitution experiment. All amplicon data was visualized using the *ggplot2* (Valero-Mora 2010).

### Stressor experiments: pathogen growth and salt tolerance

For resistance against bacterial pathogens, Ws4 and *per5* plants were grown in potting soil for 4-5 weeks. Leaves were infiltrated with a bacterial suspension *of Pseudomonas* strain *PtoDC3000/Rif* (O.D._600_=0.0001, Animal and Plant Health Inspection Agency (APHIA) import permit number: **112-B-510**) in 10mM MgSO_4_. Plants were covered to maintain high humidity. Infiltrated leaves were harvested after 3-4 days when symptoms appeared. The pathogen titer per leaf disc (8mm diameter) was quantified through a viable plate counting method after serial dilution. For salt tolerance, plants were germinated on agar plates for 7 days before being transferred to freshly prepared agar plates with the indicated concentrations of NaCl. Alternatively, plants were grown in potting soil with a regular watering scheme. Salt solution of the indicated concentration was applied in week 2 and week 3. Shoot fresh weights were measured at day 17 for agar plate-grown plants or day 35 for soil-grown plants.

### Transcriptome experiments

Plants were germinated with the *At-*16SC1 for 14 days before harvesting. Roots and shoots were harvested using the same procedure above. RNA was extracted with the Plant RNeasy Mini Kit (Qiagen) according to the manufacturer’s instructions. After a quality check, libraries were prepared by Novogene-Europe and sequenced on Illumina platform PE150 each with 3Gb sequencing depth.

Raw Illumina RNA-Seq reads were pre-processed using fastp (Chen et al. 2018) with default settings for paired-end. High quality reads were pseudo-aligned to TAIR10.58 *Arabidopsis thaliana* transcriptome reference (Ensembl)(Yates et al. 2022) using kallisto (Bray et al. 2016). On average, we obtained 11.61 million paired-end reads per sample. After removal of low abundant transcripts that were absent in at least two replicates under each condition, count data were imported using the *tximport* package (Soneson et al. 2016). The log_2_ scaled counts were normalized by the identified SVs (Leek et al. 2012) using the *limma* package (*removeBatchEffect* function) (Ritchie et al. 2015), and transformed as median-centered *z*-score by transcripts (scaled counts, *scale* function). Then *z*-scores were used to conduct *k*-means clustering for all transcripts. The cluster number (*k* = 8) was determined by sum of squared error and Akaike information criterion.

Differential expression analyses were performed using the *DESeq2* package(Love et al. 2014). Pairwise comparisons were designed as: (1) *per5* mock vs WT mock, (2) SynCom-treated WT vs WT mock, (3) SynCom-treated *per5* vs SynCom-treated WT, (4) SynCom-treated *per5* vs *per5* mock. Transcripts with fold-changes > 1.5 with adjusted *p*-value equal to or below 0.05 were considered significant.

Gene ontology (GO) enrichment for each cluster using the whole *Arabidopsis* transcriptome as background were performed with the *GOseq* package (Young et al. 2010) with the consideration of transcript length. GO annotations were retrieved from the Gene Ontology Consortium (September 2019). Significantly changed biological process GO terms (adjusted *p*-value < 0.05) were visualized in dot plots using the *clusterProfiler* package (Yu et al. 2012).

### Statistical analysis and data availability

Analyses were performed using the R environment. Dunn’s Kruskal-Wallis and ANOVA were used to test for statistical significance. A *p*-value smaller than 0.05 was considered significant unless otherwise specified. The scripts used *16S* amplicon sequencing and RNA-seq analysis are available upon request.

## Author contributions and acknowledgement

K.-W. M. was involved in the conceptualization and writing of the manuscript with inputs from all co-authors. K.-W.M. performed the initial screening, RNA-seq and *16S* community profiling. T.-T. L. performed functional characterization of the *per5* mutant including chemicals treatments and functional reconstitution experiments. M. I. analyzed the RNA-seq data. K.-W. M. and H.-J. S. performed experiments characterizing the growth phenotype of *per5* in natural soils and under salt treatment. K.-W. M. and S.P. performed the experiment quantifying the levels of SA and JA.

We thank Dr. Paul Schulze-Lefert for general discussion. We thank Dr. Pengfan Zhang for the assistance in using the rbec pipeline. This research was funded by the Deutsche Forschungsgemeinschaft (DFG, German Research Foundation) under the project number **SPP 2125 DECRyPT MA 9714/1-1** and the National Science and Technology Council (NSTC, Taiwan) under the project number **113-2311-B-001-009-MY3** to K.-W.M.

---

**Supp S1.**
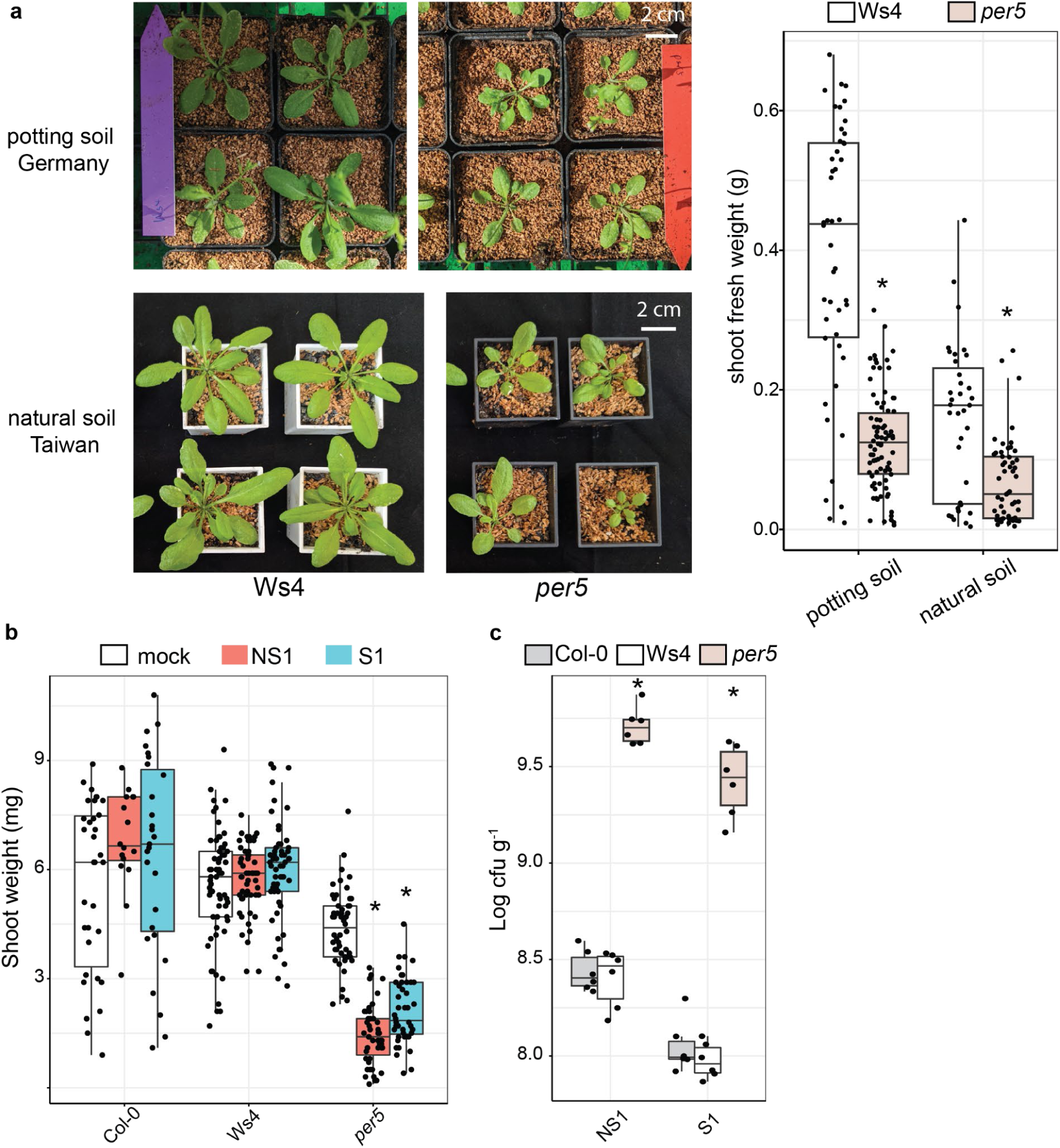
*per5* phenotypes are reproducible using synthetic communities with contrasting immune-activating and suppressive traits. (a) Representative images of 5-week old Ws4 and *per5* grown in peat-based potting soil and natural soil. Experiments were performed separately in two greenhouses in Germany and Taiwan. Shoot fresh weights of plants grown in both potting soil and natural soil collected from Taiwan. Statistical significance was determined by ANOVA. Different asterisks indicate statistical significance of *p*≤0.05 compared to the wild type Ws4 plants. (b) and (c) Shoot fresh weight and total microbial load of two-week old plants inoculated with the 5-member immune non-suppressive (NS1) and suppressive SynCom (S1). Statistical significance was determined by Kruskal-Wallis followed by Dunn’s post-hoc test adjusted by Benjamini-Hochberg method. Different asterisks indicate statistical significance of *p*≤0.05 compared to the mock control or the corresponding wild type parent unless otherwise specified.

**Supp S2.**
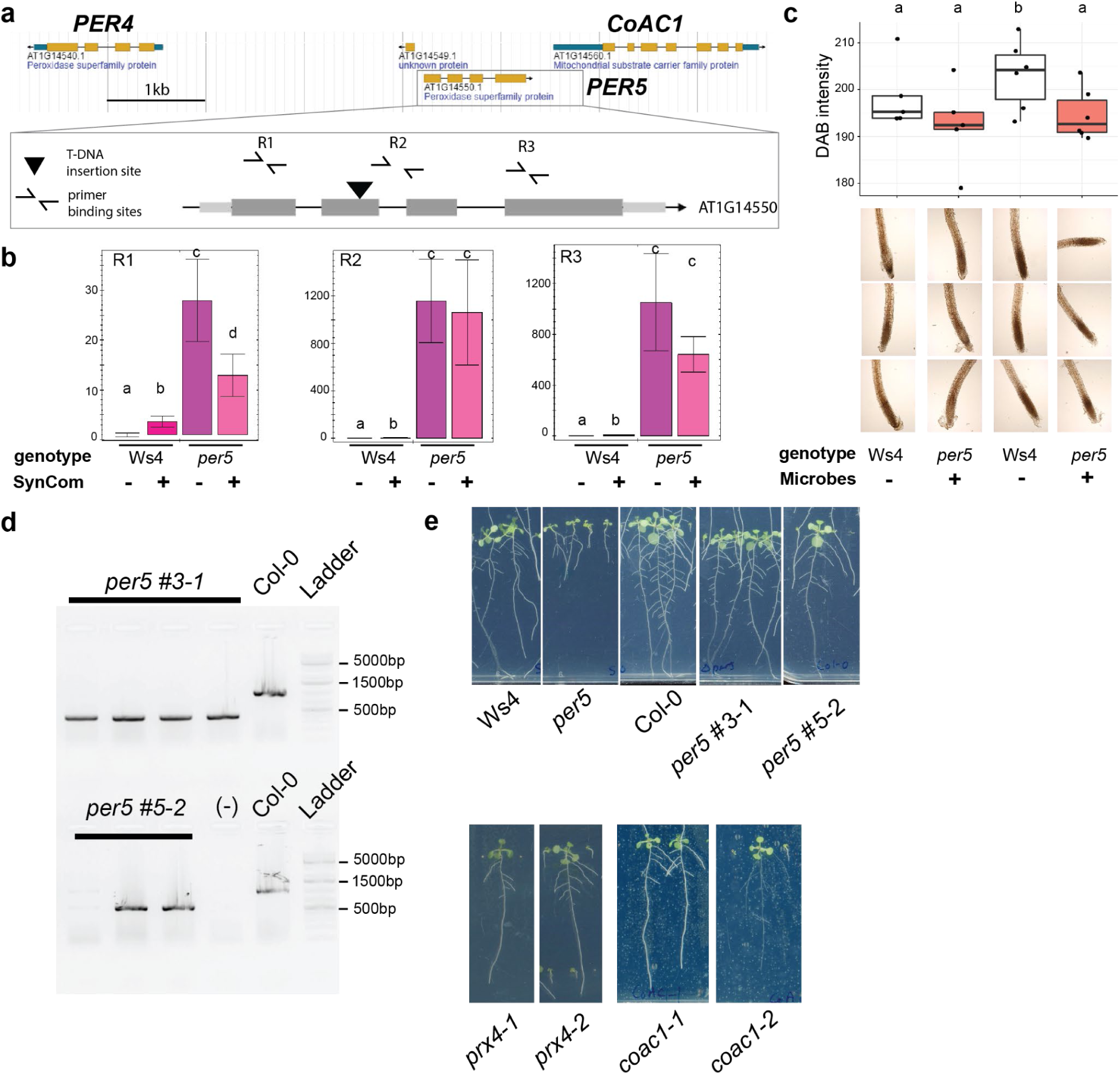
Mutation of the PER5 gene is not sufficient to cause dysbiosis phenotypes. (a) Diagram of the genetic loci centered on *PER5* (AT1G14550). A T-DNA (indicated by a block triangle) is inserted into the second exon of *PER5* mutant line FLAG_264G01. (b) The expressions of transcripts upstream and downstream of the T-DNA insertion site as quantified by quantitative RT-PCR. (c) Intensity of DAB as a proxy of the level of reactive oxygen species under mock treatment or bacterial inoculation. Plants are inoculated with the *Pseudomonas* strain R9. (d) Confirmation of deletions in two independent *PER5* CRISPR lines by PCR. Accession Col-0 was included as a control. (e) Representative images of *PER5* deletion mutants (per5 #line 3-1 and 5-2), *PER4* knock-down mutant (SALK_110617C, *prx4-1*), PER4 hypermorphic mutant (SALK_044730C, *prx4-2*) and *COAC1* null mutants (SALK_091671C, *coac1-1*; SALK_087365C, *coac1-2*) two weeks after inoculation with the 16-member SynCom. Statistical significance was determined by ANOVA. Different letters indicate statistical significance of *p*≤0.05.

**Supp S3.**
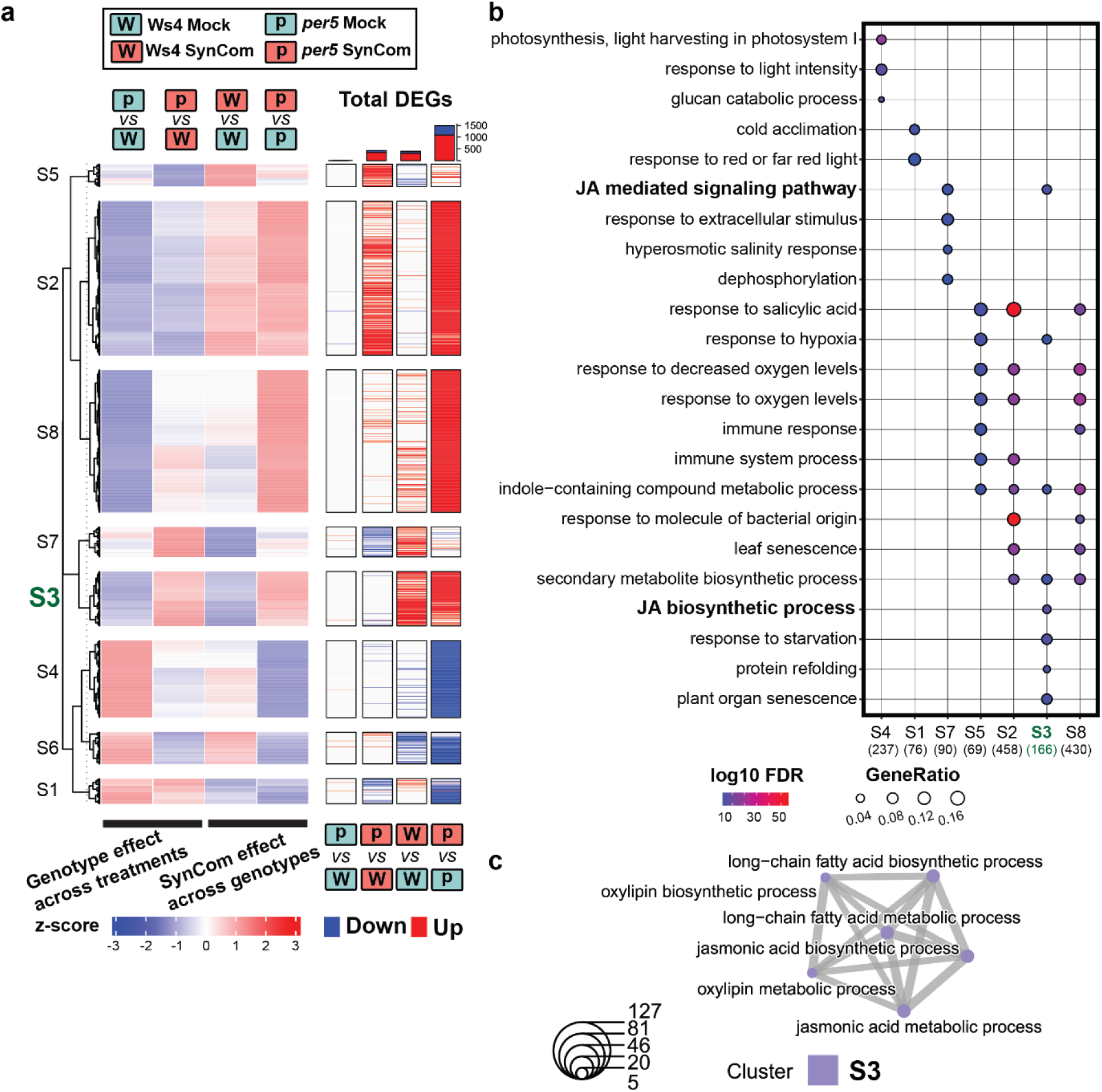
Shoot transcriptomes of WT and *per5* plants in response to SynCom. (a) Heatmaps summarizing the relative expressions of genes across genotypes in shoots. DEGs were calculated by pairwise comparisons (log2FC>1.5, *p*<0.05). (b) Significantly enriched GO terms associated with each cluster for the shoot dataset. (c) GO terms related to JA processes and their association with clusters S3. Size of pie charts corresponds to the number of DEGs.

**Supp S4.**
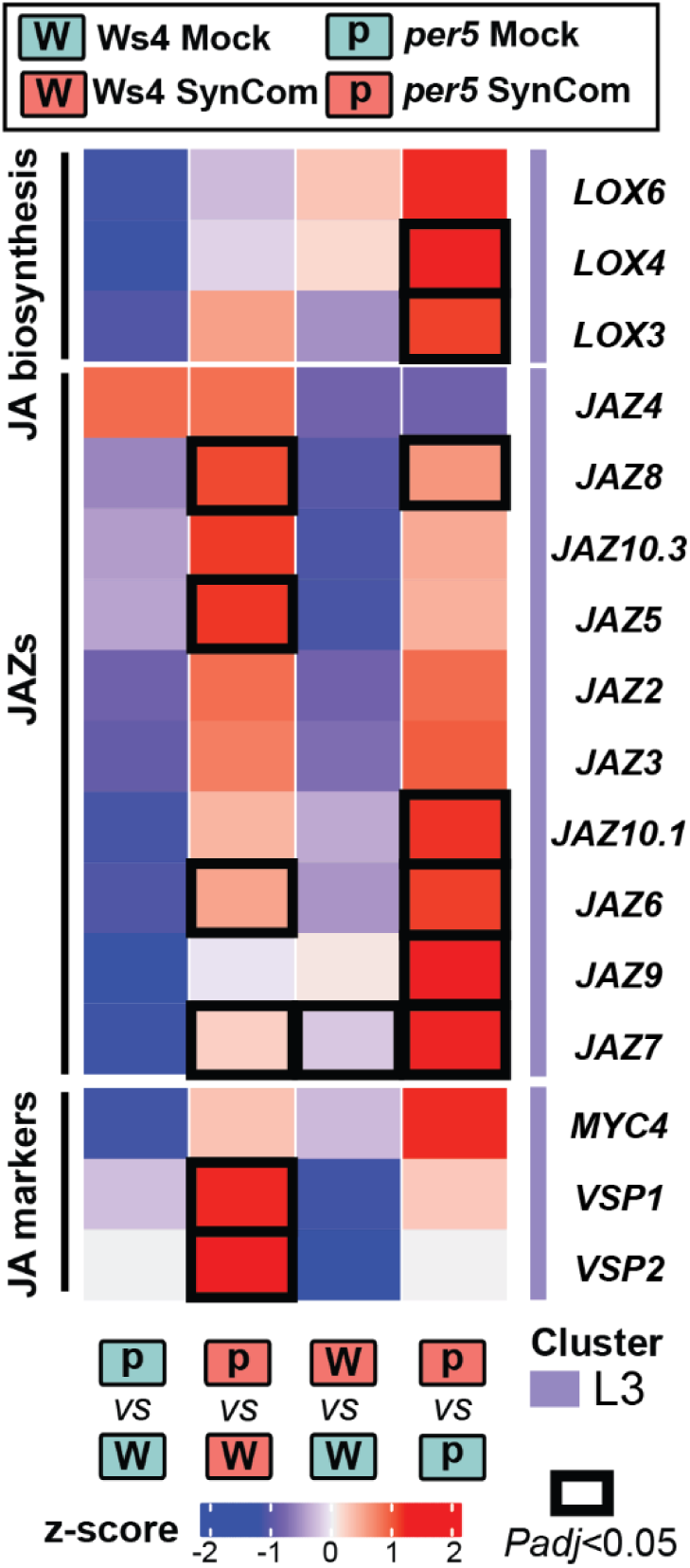
JA related genes are upregulated in *per5* in response to the synthetic community in shoot. Normalized relative expression levels of selected genes involved in JA-related processes in shoots (related to Fig 3b). Genes of statistical significance (log2FC>1.5, *p*<0.05) are indicated with blocked lines.

**Supp S5.**
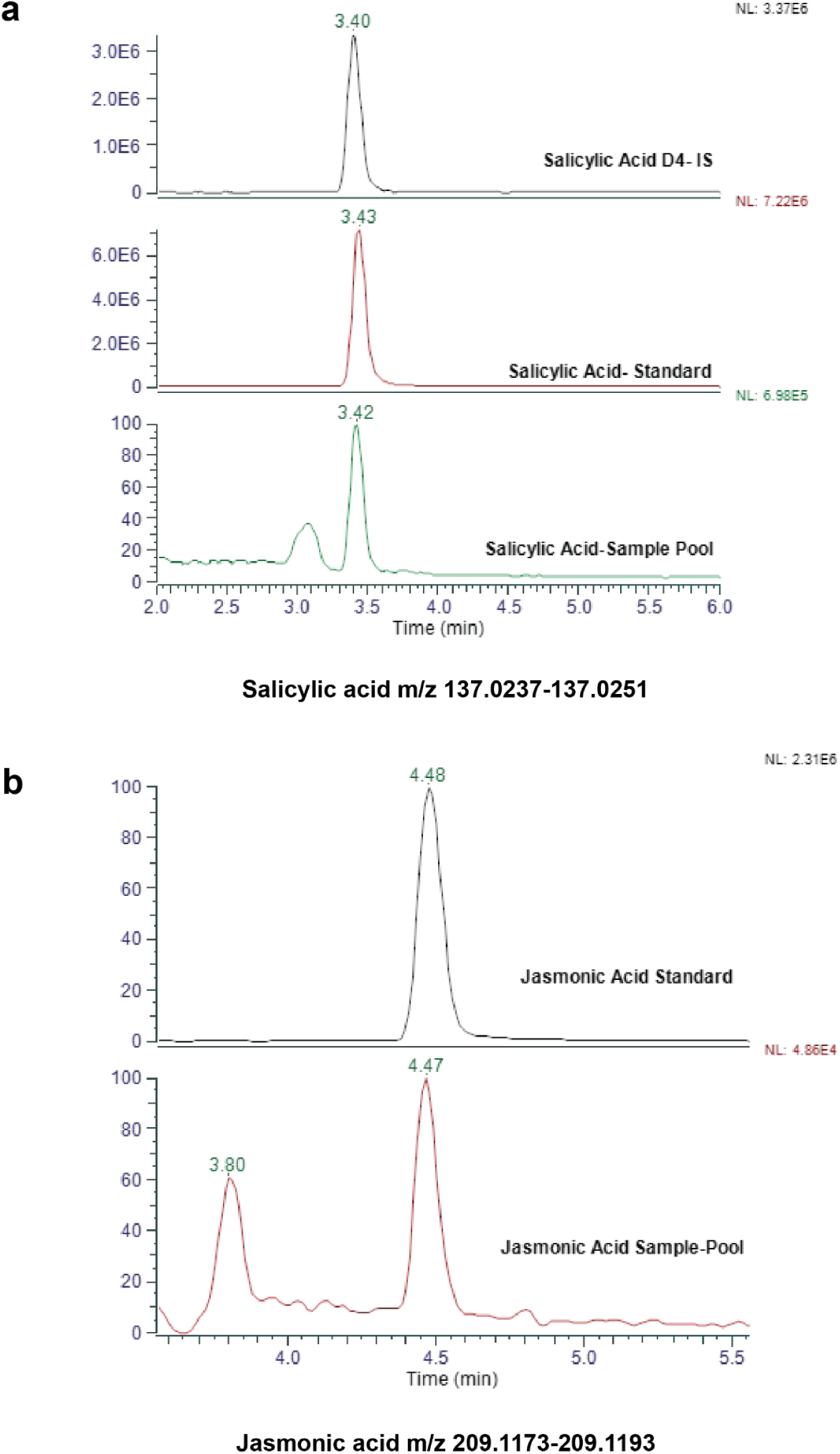
Quantification of JA and SA for WT and *per5* plants. Overlay chromatograms of JA and SA with their corresponding internal standards.

